# Characterization of methods for mechanistic inference of the gut microbiome in disease

**DOI:** 10.1101/2023.12.01.569617

**Authors:** Brook Santangelo, Lawrence Hunter, Catherine Lozupone

## Abstract

**Motivation:** Knowledge graphs have found broad biomedical applications, providing useful representations of complex knowledge. Although plentiful evidence exists linking the gut microbiome to disease, mechanistic understanding of those relationships remains generally elusive. A structured analysis of existing resources is necessary to characterize the resources and methodologies needed to facilitate mechanistic inference.

**Results:** Here we demonstrate the potential of knowledge graphs to hypothesize plausible mechanistic accounts of host-microbe interactions in disease and define the need for semantic constraint in doing so. We constructed a knowledge graph of linked microbes, genes and metabolites called MGMLink, and one of microbial traits, environments, and human phenotypes called KG-microbe-phenio. Using a shortest path search and a pattern based semantically constrained path search through the graphs, we highlight the need for a microbiome-disease resource and semantically informed search methods to enable mechanistic inference.

**Availability:** The software to create MGMLink is openly available at https://github.com/bsantan/MGMLink, and KG-microbe is available at https://github.com/Knowledge-Graph-Hub/kg-microbe and KG-phenio is available at https://github.com/Knowledge-Graph-Hub/kg-phenio.

## 1 Introduction

That gut microbiome composition differs in disease states has become increasingly clear from metagenomic studies (Falony et al. 2019; King et al. 2019). However, our understanding of specific microbiome signatures on clinical outcomes is sparse and mainly correlative (Huang et al. 2017; Chang and Kao 2019). Microbiome signatures have been associated with auto-immune, gastrointestinal, neurological disease and cancer (Berg et al. 2020; Stopińska, Radziwoń-Zaleska, and Domitrz 2021), but the conclusions often lack a mechanistic account. Experimental studies to test hypothetical mechanisms are critical, but they are expensive and cannot cover all potential pathways a microbe might act on within a host. Additionally, investigations of immune and metabolomic outcomes in relation to microbial abundance lack a comprehensive integration of biomedical knowledge. Without this mechanistic account we do not fully understand the role of the gut microbiome in disease, which makes the development of targeted therapeutics more difficult.

With more than 20,000 citations relating the microbiome to disease published each year in PubMed since 2019, there is huge potential for integrating existing studies into a centralized knowledge base (Badal et al. 2019). Understanding mechanisms by which the gut microbiome may impact disease requires a comprehensive view of how microbes and their metabolic products interact with the human body. We therefore seek to provide contextual insight to predict how microbes influence disease.

To address the limitations of correlative analysis, we present a methodology that integrates microbiome information into knowledge graphs (KGs) and generates hypothetical mechanistic accounts of how microbes influence disease. KGs describe relationships between entities of different types, *e*.*g*., how a gene or gene product influences a disease, or how a metabolite is processed in a specific pathway. KGs have broad applications across biomedical research, including prediction of drug-drug interactions (Nicholson and Greene 2020; Reese et al. 2021; Tripodi et al. 2020), evaluation of mechanisms of toxicity (Tripodi et al. 2020), prediction of unknown drug disease targets (Mayers et al. 2022), or linking symptoms from electronic health records to better understand disease (Zhang et al. 2019). However, these applications have yet to be extended into the microbiome field (Zhang et al. 2019; Badal et al. 2019). Existing microbial KGs incorporate information more focused on microbial trait outcomes (Joachimiak et al. 2021) or are limited in scope and/or lacking in biomedical information (King et al. 2019; Liu et al. 2020; Ma et al. 2017). Here we describe the creation and application of two KGs of microbial and biomedical content in the successful, or unsuccessful, creation of mechanistic hypotheses regarding microbe-disease relationships, and propose requirements for future resources and methodologies to construct these hypotheses.

We created a microbiome relevant KG representing microbe, gene, metabolite, disease links called MGMLink by integrating known microbe-host interactions into a biomedical KG built using the PheKnowLator system (Figure 1). PheKnowLator is a Python 3 library that enables construction of KGs that incorporate a wide variety of data and terminology sources, including ontologies such as the Monarch Disease Ontology (MONDO), the Chemicals Entities of Biological Interest Ontology (CHEBI), the Gene Ontology (GO) and the Human Phenotype Ontology (HPO) (Callahan et al. 2020). To augment the default PheKnowLator KG with information on microbes, we integrated data from gutMGene, a manually curated repository of assertions involving gut microbes, microbial metabolites, and target genes from over 360 PubMed publications (Cheng et al. 2022). We also created a microbiome-relevant KG, KG-microbe-phenio, by integrating the microbe-oriented KG-microbe with the phenotype-oriented KG-phenio. KG-microbe was built from a resource of phenotypic microbial traits by Madin et al., and relevant ontologies including CHEBI, the Environment Ontology (ENVO), GO, and NCBITaxon (Joachimiak et al. 2021). KG-phenio integrates Monarch ontologies including MONDO, HPO, and GO allowing for human disease content to be introduced to KG-microbe (https://github.com/Knowledge-Graph-Hub/kg-phenio). Each KG therefore integrates a range of relevant ontologies (Table 1). Using MGMLink and KG-microbe-phenio, we examine mechanisms that describe the interaction between a microbe and a disease. We explore the extent to which previously identified microbes influencing mental disorders could be further examined with the structured content of these KGs. We applied structure and semantic based algorithms to assess the mechanistic feasibility of resulting paths. Our results indicate that (1) current resources do not adequately depict microbial function and (2) data-driven approaches based on multi-omic studies can greatly improve the mechanistic feasibility of paths found. This analysis presents evidence that KGs have potential in solving the important problem of mechanistic inference in the field of the gut microbiome and suggests specific gaps in the resources and methodologies to perform this task.

**Table 1.** Categories of nodes that exist MGMLink and KG-microbe-phenio and are used to characterize path patterns.

| Description | Resource/Ontology | Example Node Types |
| --- | --- | --- |
| Metabolite, chemical class | Chemical Entities of Biological Interest (CHEBI) | Pyruvate, nitrogen dioxide |
| Protein | Protein Ontology (PRO) | Apoptosis regulator BAX (human), tryptophan 5-hydroxylase 1 (human) |
| Disease, disorder | Monarch Disease Ontology (MONDO) | Depressive disorder, cognitive disorder |
| Gene | NCBI gene (gene) | TPH1 (human), BAX |
| Organism | NCBI Taxonomy (NCBITaxon) | Lactobacillus plantarum, Bacteroides |
| Organism in an anatomical context | NCBI Taxonomy, UBERON (Contextual Microbe) | Lactobacillus plantarum in a human intestine |
| Molecular function, biological process, cellular component | Gene Ontology (GO) | Cyclic nucleotide-mediate signaling, serotonin biosynthetic process |
| Phenotype | Human Phenotype Ontology (HPO) | Tremor, Abnormality of the cardiovascular system |
| Anatomical entity | Uber-Anatomy Ontology (UBERON) | Intestine, brain |
| Supporting information retrieval | Basic Formal Ontology (BFO) | Role, process |
| Anatomical term across species | Common Anatomy Reference Ontology (CARO) | Biological entity, anatomical entity |
| Environmental system, component, process | Environment Ontology (ENVO) | Acidic environment, fecal environment |
| Ecological trait, organism | Ontology of Core Ecological Entities (ECOCORE) | Aerobe, facultative aerobe |
| Phenotype across species | Unified Phenotype Ontology (UPHENO) | abnormality of embryonic tissue physiology, abnormal phenotype by ontology source |
| Mammalian phenotype | Mammalian Phenotype Ontology (MPO) | abnormal serotonin level, abnormal nervous system dopamine level |
| Cell line | Cell Line Ontology (CLO) | Platelet, T cell |

**Figure 1.**
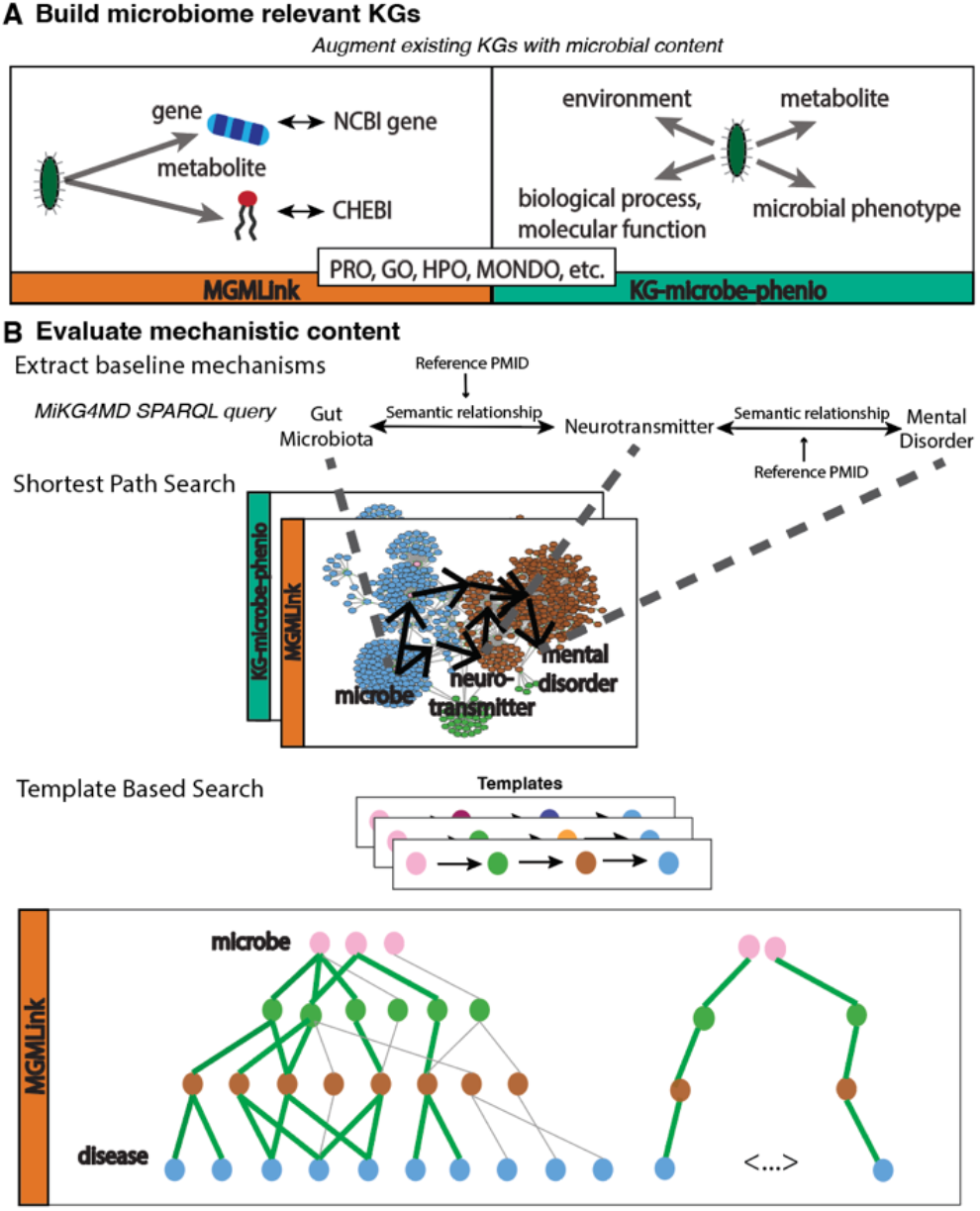
Methodological approach. (A) Microbial content was introduced to the biomedical KG generated using PheKnowLator from GutMGene, and to the biomedical KG-phenio from KG-microbe. (B) We evaluated microbe-neurotransmitter-disease pairs based on what was found in the MiKG4MD resource using a shortest path search and a template-based search.

## 2 Methods

### 2.1 MGMLink Construction

To generate MGMLink, we incorporated previously published microbe-host interactions from the gutMGene database into the PheKnowLator framework (Figure 1A). The framework allows alternative knowledge modeling approaches; we used a model in which concepts are unidirectionally relationally linked in the graph (as opposed to bidirectionally) (Callahan et al. 2020). We also use a version of a PheKnowLator KG that reflects a transformation based on OWL-NETS, which enables network inference of OWL-encoded knowledge via abstraction into biologically meaningful triples (Callahan et al. 2018). These parameters produce the topologically simplest graph in the most performant manner in relevant metrics such as node embedding quality. The representation of gutMGene data in the framework was matched to the PheKnowLator framework parameters. All assertions were mapped to an OWL-encoded KG representation and then transformed into triples using OWL-NETS, resulting in four unique patterns, examples of which are shown in Table 1 (Callahan et al. 2018). This resulted in new KG nodes that represented microbes in the context of the anatomical location (*i*.*e*., the gut) and species (human or mouse) in which the interactions have been reported to occur.

The gutMGene database consists of microbe-metabolite and microbe-gene assertions that occur in the host of either a human or mouse. These relationships were manually extracted from over 360 PubMed publications and are based on validated methods such as RT-qPCR, high-performance liquid chromatography, and 16S rRNA sequencing (Cheng et al. 2022). Specifically, the gutMGene database describes four different types of assertions: (1) microbial substrates observed in humans or mice, (2) microbial metabolites observed in humans or mice, (3) genes observed to have negatively associated expression with microbes in humans and mice, and (4) genes observed to have positively associated expression with microbes in humans and mice (Cheng et al. 2022). Assertions 1 and 2 were extracted from the Association between Gut microbe and Metabolite v1.0 results for Human and Mouse, respectively, and assertions 3 and 4 were extracted from the Association between Gut microbe and Gene v1.0 for Human and Mouse, respectively. These were then represented using a specific semantic pattern to encompass the relationship type and the context (Table 2). We attempted to map each microbial type to an entry in the NCBI Taxonomy, and for those microbial types that could not be mapped to NCBI Taxonomy entries, new nodes representing unclassified bacterial organisms were created. Additionally, microbes that had an unclear or indirect mapping to a NCBITaxon identifier were corrected. A total of 1874 assertions from gutMGene were created in MGMLink for 533 unique microbial taxa (at the family, genus, species, or strain level). NCBI Taxonomy entries were found for 336 of these taxa, and gene or metabolic information is connected to 461 of the gutMGene assertions. To generate a connected graph, species or strains were related to their corresponding higher-level classifications (genus, family, class, etc.) which already existed in the KG. This allowed for inferences to be made regarding species-strain or genus-species relationships.

**Table 2.** Specific examples of all types of patterns identified from gutMGene assertions, and corresponding OWL-NETS representations into which these assertions were converted to integrate them into the knowledge base.

| New KG Assertion | OWL – NETS Triple Representation |
| --- | --- |
| *Streptococcus* in lower digestive tract in human inhibits CCL20 | St subclass_of NCBITaxon: Streptococcus<br>St located_in UBERON: 'lower digestive tract'<br>St located_in NCBITaxon: 'Homo sapiens'<br>St indirectly_negatively_regulates_activity_of NCBIgene: CCL20 |
| *A. muciniphila* in lower digestive tract in human activates IL10 | Am subclass_of NCBITaxon: 'Akkermansia muciniphila'<br>Am located_in UBERON: 'lower digestive tract'<br>Am located_in NCBITaxon: 'Homo sapiens'<br>Am indirectly_positively_regulates_activity_of NCBIgene: IL10 |
| *F. prauznitzii* in lower digestive tract in mouse produces benzoic acid | Fp subclass_of NCBITaxon: 'Faecalibacterium prauznitzii'<br>Fp located_in UBERON: 'lower digestive tract'<br>Fp located_in NCBITaxon: 'Mus musculus'<br>Fp contains_process o has_output CHEBI: benzoic acid |
| *R. bromii* in lower digestive tract in human metabolizes propionate | Rb subclass_of NCBITaxon: 'Ruminococcus bromii'<br>Rb located_in UBERON: 'lower digestive tract'<br>Rb located_in NCBITaxon: 'Homo sapiens'<br>Rb involved_in_positive_regulation_of o has_primary_input_or_output CHEBI: propionate |

### 2.2 KG-microbe-phenio Construction

The second resource is a biomedically relevant KG of microbial traits that we will refer to as KG-microbe-phenio. KG-microbe consists of phenotypic traits and environmental descriptors of microbes in all environments, not specifically the human gut (Joachimiak et al. 2021) (Figure 1B). The KG is a comprehensive data source that synthesizes microbial phenotypic traits, quantitative genomic traits, and habitats by employing named entity recognition and natural language processing techniques to normalize terms according to ontologies (CHEBI, GO, UBERON, etc). The phenotypic and trait information is derived from a unified dataset of merged content from GenBank, Bergey’s Manual of Systematics of Archaea and Bacteria, and literature (Madin et al. 2020). The Environment Ontology (ENVO), CHEBI, GO, and NCBITaxon are also introduced to the trait information. To incorporate molecular and human content, we combined KG-phenio, which consists of relationships among phenotypes, processes, and disease from ontologies including HPO, GO, CHEBI, and MONDO, with KG-microbe such that human phenotype information would be connected, using the Knowledge Graph Exchange (KGX) Python library (“GitHub - Monarch-Initiative/phenio” n.d.).

### 2.3 Mechanism Prediction Framework

The path between two nodes in a KG can be drawn in a nearly infinite number of ways, especially for KGs of this scale (over 780,000 nodes and 5,000,000 edges in MGMLink and over 500,000 nodes and 1,000,000 edges in KG-microbe-phenio). Here we employ a shortest-path search using a breadth-first search algorithm with unweighted edges, which incrementally searches all neighbors of a source node until the target node is found and returns the path with the minimum number of edges. Shortest path search is a common approach to examining a mechanistic link between two nodes (Zhao et al. 2020; Dijkstra 1959). The result is unbiased, allowing for a comprehensive assessment of potential interactions between microbes and metabolites, proteins, or processes of interest. We excluded specific edge types, e.g., “only_in_taxon” and “part_of” in MGMLink and “biolink:category” and “biolink:in_taxon” in KG-microbe-phenio, to remove paths irrelevant to the mechanistic accounts we are seeking.

We evaluated the content of MGMLink and KG-microbe-phenio by searching for pairs identified in a previously curated microbiome mental disorder based KG called MiKG4MD (Liu et al. 2021). MiKG4MD was built from manually annotated relationships from literature including 6 neurotransmitters, 10 mental disorders, 45 microbes, and 56 KEGG pathways. All connected triples consisting of (1) microbe, related to, mental disorder or (2) microbe, increases/decreases, neurotransmitter and neurotransmitter, related to, mental disorder were found in MiKG4MD using SPARQL queries. We characterized the content of each shortest path by categorizing each node type within the path among all ontologies that exist in the KG (Table 1, Figure 1B). The result is a set of patterns which describe the composition of all paths from the shortest paths search. Each pattern consists of the unique types of nodes among paths.

We next evaluated the content of MGMLink, which includes more molecularly plausible paths consisting of the microbe-host interactions represented in the KG. We applied a template-based search, using the template *microbe, metabolite, gene, protein, process, disease*, over MGMLink paths (Figure 1B). The path followed in this template includes microbial metabolites that affect human gene expression (*microbe, metabolite, gene*) and how those gene products functionally are implicated in disease (*protein, process, disease*). We performed a bidirectional search within the graphs for connected nodes that follow these patterns (*e*.*g*., microbe - gene and gene - protein), and constructed paths with all combined triples. We evaluated these paths for common occurrences of metabolite-disease pairs.

## 3 Results

### 3.1 Shortest Path Search

To assess the content and quality of MGMLink and KG-microbe-phenio, we evaluated the existence of node types within the shortest path between each microbe-disease pair, and each microbe-neurotransmitter and neurotransmitter-disease combined pair. By ignoring the order of node types or abundance of each node type within the path, this evaluation simply characterized the edge and node types among each KG. When comparing paths between microbe-disease pairs, paths which included metabolites (CHEBI nodes) were more common in MGMLink, with the most common pattern including microbes, metabolites, and diseases (*microbe-chebi-mondo)*. An example path with this pattern has some mechanistic feasibility, as 3-indolepropionic acid (IPA), a metabolite of tryptophan, is associated with neuroinflammation, however ketoprofen and IPA are connected through a common parent chemical, propionic acid, making the mechanistic qualities less concrete and specific (Figure 2A) (Konopelski and Mogilnicka 2022). Alternatively, paths from KG-microbe-phenio most often include microbes and diseases (*microbe-mondo)* or microbes, anatomical entities, and diseases (*microbe-mondoenvo-uberon)*, patterns which commonly have paths going through several unrelated diseases and very broad terms such as “disease or disorder” or unrelated environments including “mouth environment”, “brain”, or “respiratory tract” (Figure 2A). Results such as this indicate nodes and edges to avoid, or patterns that arrive at high level concepts, such as multiple diseases that are subclasses of “disease or disorder”. In both KGs, the neurotransmitter of interest is absent from all paths, prompting the need for a more targeted search.

**Figure 2.**
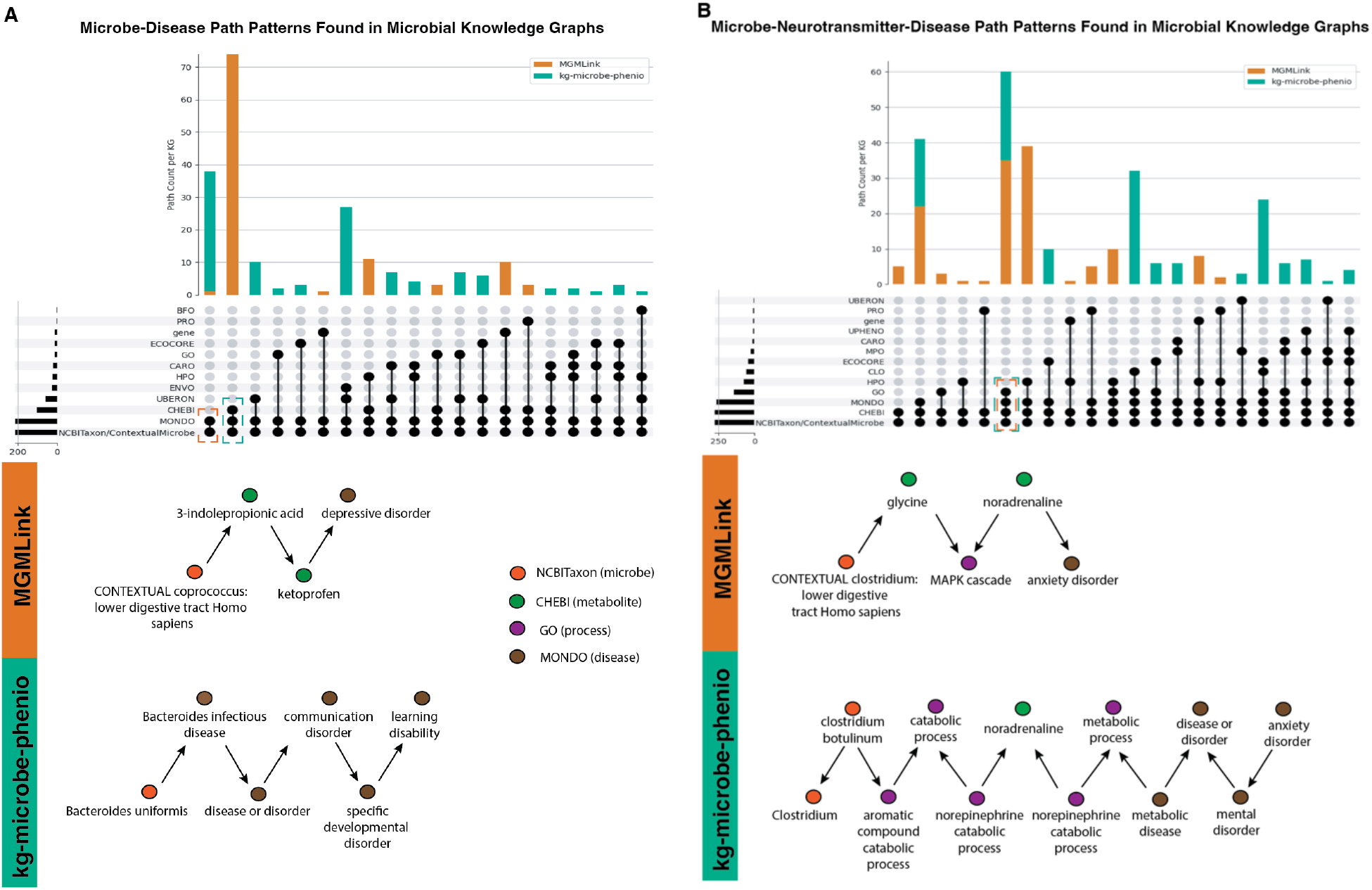
Analysis of existing microbial knowledge bases. (A) Patterns for shortest path search from each microbe to each disease as found in MiKG4MD. Examples of most common patterns for each KG are shown below. (B) Patterns for shortest path search from each microbe to each neurotransmitter, then that neurotransmitter to the corresponding disease as found in MiKG4MD. Examples of one commonly found pattern in both KGs are shown below.

When the intermediate neurotransmitters are added to the path search, the patterns include phenotypes or GO processes more often. MGMLink has the two most common patterns of microbes, metabolites, phenotypes, and diseases (*microbe-chebi-mondo-hpo)* and microbes, metabolites, processes or biological functions, and diseases (*microbe-chebi-mondo-go)*. This allows for more detailed characterization of microbial influence in the host by including host attributes, such as the MAPK cascade which has been implicated in anxiety (Figure 2B) (Wefers et al. 2012). KG-microbe-phenio similarly has a common pattern of paths including microbes, metabolites, processes or biological functions, and diseases (*microbe-chebi-mondo-go)*, however path are much longer, and concepts are much broader. The path shown includes “catabolic process”, “metabolic process”, and “disease or disorder”, all of which are too broad to gain mechanistic insight (Figure 2B). This investigation allowed us to identify specific qualities of mechanistic or non-mechanistic paths and exemplified the need for a more effective path search methodology for microbiome research. Due to these limitations in current representations of microbial knowledge, it is necessary to develop a comprehensive resource consisting of knowledge from a range of microbial databases and ontologies.

### 3.2 Template Based Approach

Next, we applied a template-based approach to this path search through MGMLink to evaluate whether more plausible mechanisms might be uncovered. The resulting template-based paths that connected microbes with diseases were enriched in specific metabolites, which suggests common aspects of this microbial interaction in a disease. Genistein and quercetin are metabolites that most often were part of the *microbe, metabolite, gene, protein, process, disease* paths (Figure 3). Quercetin has been cited as a prebiotic with anti-inflammatory and neuroprotective effects, but the diseases shown to be associated have no clear pattern (Balasubramanian et al. 2022). Choline is another metabolite that is commonly found across many diseases and has been implicated in multiple metabolic disorders, and folic acid is a known microbial metabolite involved in biochemical processes, gene regulation, and intestinal integrity (Spencer et al. 2011). To further validate whether trends uncovered by this search can fully depict mechanistic paths, we investigated a specific example of a path including folic acid. The path infers that the inflammatory state affected by folic acid may be through IL-18, which is a proinflammatory cytokine, which is mechanistically feasible (Hossain, Amarasena, and Mayengbam 2022). The direction of this interaction, however, is not depicted here and further literature investigation would be required to understand that folic acid is anti-inflammatory, and negatively correlated with IL-18 and inflammation.

**Figure 3.**
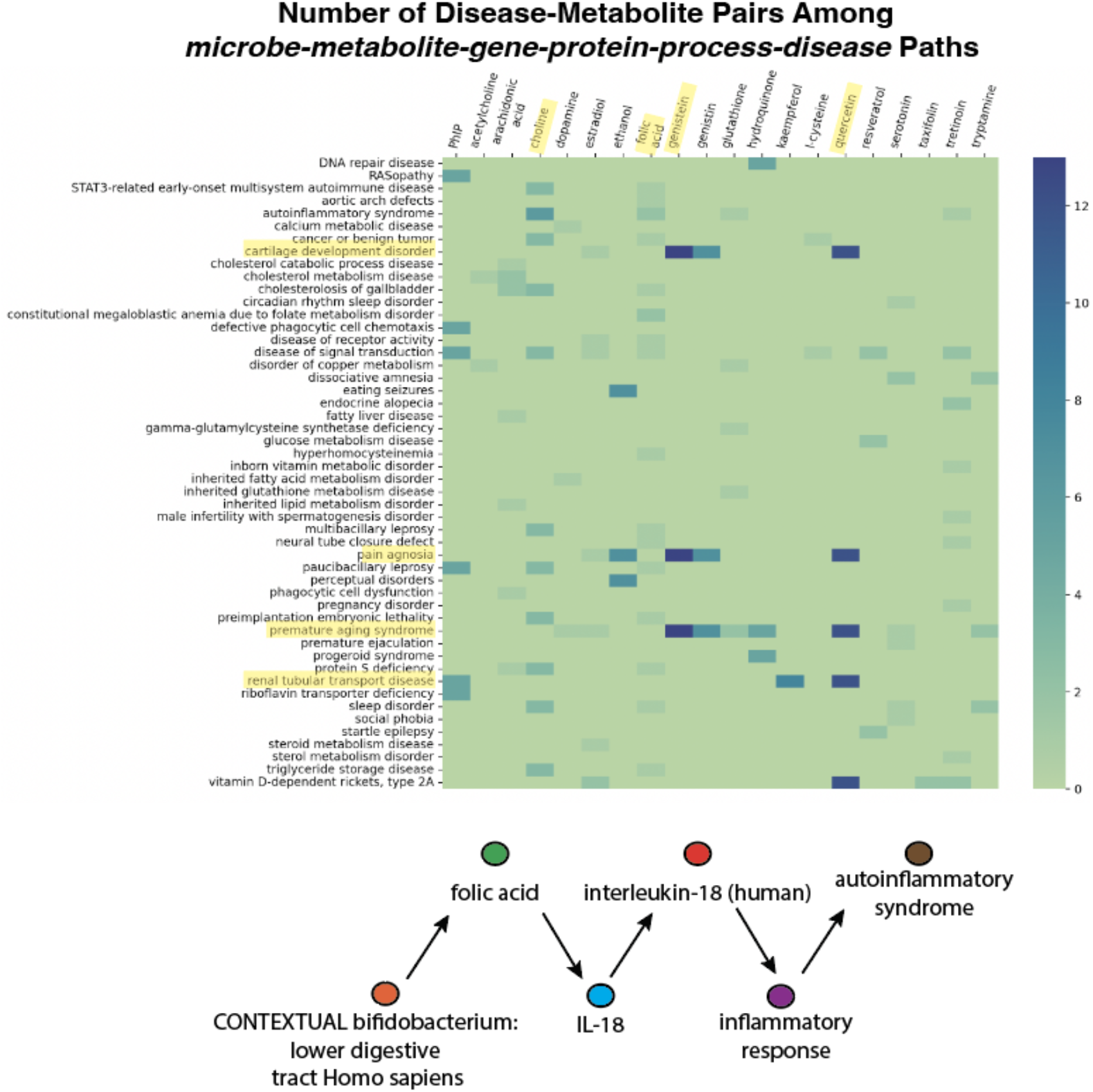
Number of disease-metabolite pairs that exist for all paths found that follow the microbe, metabolite, gene, protein, process, disease template. Highlighted diseases and metabolites show most common instances. An example of a path containing folic acid is shown below.

## 5 Discussion and Conclusion

There are many resources available that represent microbial metabolism, pathways, genomic content, and taxonomy as well as attempts to integrate this information in a manual or automated way. Databases of reference genomes include the NCBI Reference Sequence Database (RefSeq), the NIH genetic sequence database (GenBank) and the International Nucleotide Sequence Database Collaboration (INSDC), from which standardized taxonomic databases such as NCBI Taxonomy (NCBITaxon) and the Genome Taxonomy Database (GTDB) classify organisms and standardize naming conventions (Pruitt, Tatusova, and Maglott 2007; Sayers et al. 2022; Arita, Karsch-Mizrachi, and Cochrane 2021; Federhen 2012; McDonald et al. 2018). These organism centric databases are referenced in pathway databases including Reactome, Meta-Cyc, BiGG, and KEGG, all of which represent genome scale metabolic network models (GSMNs) or pathway/genome databases (PGDBs) (Gillespie et al. 2022; Caspi et al. 2020; Norsigian et al. 2020; Kanehisa and Goto 2000). Several microbial databases exist which infer the metabolic pathways and functional potential of microbial genomes including the MetAboliC pAthways DAtabase for Microbial taxonomic groups (MACADAM), the Microbial Signal Transduction Database (MIST), and AnnoTree, that vary in applications from functional inference to metabolic signal transduction to evolutionary hypothesis generation, respectively (Le Boulch et al. 2019; Gumerov et al. 2020; Mendler et al. 2019). MetaNetX is one resource that cross-links GSMNs to merge reactions, metabolites, enzymes, and pathways from the above resources with common mappings (Moretti et al. 2021). The Virtual Metabolic Human database (VMH) similarly integrates GSMNs and performs metabolic reconstructions with the goal of inferring biomedical interactions among metabolism, disease, nutrition, and the microbiome (Noronha et al. 2019). The Human Metabolome Database (HMDB) consists of small molecule metabolites found in the human body and provides mappings to pathways (KEGG), proteins (UniProt, the Protein Data Bank), and chemical databases (PubChem, CHEBI) (Wishart et al. 2022; Berman, n.d.; UniProt Consortium 2019). The Human Microbial Metabolome Database (MiMeDB) is yet another multi-omic resource of microbial genomes and metabolites and the human exposome in how they influence human health and disease (Wishart et al. 2023).

Knowledge graphs have extensive applications in the biomedical field because they integrate complex concepts in a systematic way. Several KGs exist that automate the search and integration of databases that store microbial function and pathways for modeling of complex biological systems. BioChem4j is a KG that connects NCBI Taxonomy, UniProt, and MetaNetx using CHEBI, Rhea, and NCBITaxon identifiers (Swainston et al. 2017). UniFunc similarly automatically collects data from KEGG, MetaCyc, UniProt, and the HMDB to capture genome-scale metabolic models for specific organisms (Queirós et al. 2021). Both of these extensive resources could be further integrated with human disease and phenotype content from HPO or MONDO to enable inference of the microbiome and disease. We presented two examples of KGs that encompass aspects of the above microbial and human content, MGMLink and KG-microbe-phenio, as well as disease. MGMLink contains only the limited number of microbes described in the gutMGene database and does not cover all mechanisms by which microbes influence host genes or consume or produce metabolites. While KG-microbe-phenio has a much larger number of microbes and traits, the path types connecting traits to human processes and disease rarely include content necessary to derive a molecular mechanism as shown by the broad concepts found using shortest path search. An extensive resource that adequately captures the abundant information surrounding microbial and human metabolism, maps all identifiers, and is universally available to computationally biologists and microbiologists alike is necessary.

The shortest path search methodology allowed us to investigate the content of these resources, and the template-based approach applied to MGMLink took one step further in identifying a semantic-based methodology for mechanistic inference. Although we only discuss one template in this work which may not have adequately identified mechanistic paths in the KG, the approach can be expanded to other patterns rooted in an understanding of microbe-host interactions. With these results we propose that novel resources integrate microbial metabolism and function with human phenotypes and disease are necessary to approach this problem. Furthermore, path search methods through microbiome relevant KGs will require semantics and multi-omic results to better filter out unspecific concepts, prioritize experimentally validated relationships and interactions, and ultimately move us closer towards understanding the mechanisms of the gut microbiome in disease in an automated and efficient fashion.

## Acknowledgements

We gratefully acknowledge members of the Translational and Integrative Sciences Lab and the Lawrence Berkeley National Laboratory for their input.

